# Estimating smooth and sparse neural receptive fields with a flexible spline basis

**DOI:** 10.1101/2021.03.31.437831

**Authors:** Ziwei Huang, Yanli Ran, Thomas Euler, Philipp Berens

## Abstract

Spatio-temporal receptive field (STRF) models are frequently used to approximate the computation implemented by a sensory neuron. Typically, such STRFs are assumed to be smooth and sparse. Current state-of-the-art approaches for estimating STRFs based empirical Bayes estimation encode such prior knowledge into a prior covariance matrix, whose hyperparameters are learned from the data, and thus provide STRF estimates with the desired properties even with little or noisy data. However, empirical Bayes methods are often not computationally efficient in high-dimensional settings, as encountered in sensory neuroscience. Here we pursued an alternative approach and encode prior knowledge for estimation of STRFs by choosing a set of basis function with the desired properties: a natural cubic spline basis. Our method is computationally efficient, and can be easily applied to Linear-Gaussian and Linear-Nonlinear-Poisson models as well as more complicated Linear-Nonlinear-Linear-Nonlinear cascade model or spike-triggered clustering methods. We compared the performance of spline-based methods to no-spline ones on simulated and experimental data, showing that spline-based methods consistently outperformed the no-spline versions. We provide a Python toolbox for all suggested methods (https://github.com/berenslab/RFEst/).

## 1 Introduction

Spatiotemporal receptive fields (STRFs) are frequently used in neuroscience to approximate the computation implemented by a sensory neuron. Such models consist typically of one or more linear filters summing up sensory inputs across time and space, followed by a static nonlinearity and a probabilistic output process^1^. For this, different distributions can be used depending on the type of recorded neuronal responses.

The simplest way to estimate the STRF is arguably to compute the spike-triggered average (STA), the average over all stimuli preceding a spike^2^. However, the results of spike-triggered analysis are often noisy due to limited data and prone to be corrupted when the neuron is stimulated with correlated noise or natural stimuli^3^. The maximum likelihood estimate (MLE), also known as whitened-STA (wSTA), does not suffer from such biases, but it requires even more data to converge, making it a less preferred option in many experimental settings.

It has been suggested that these shortcomings can be overcome by computing the maximum a posteriori (MAP) estimates within the framework of Bayesian inference^4^. Under this framework, prior knowledge about STRFs such as sparsity, smoothness^5^, or locality^6^ can be encoded into a prior covariance matrix, whose hyperparameters are learnt separately from the data via evidence optimization. These algorithms thus provide STRF estimates with the desired properties even with little or noisy data. Although elegant in theory, the evidence optimization step of this framework is computationally costly: the cubic complexity in the dimensionality of the parameter space renders it difficult to scale to high-dimensional setting as often encountered in sensory neuroscience. Also, current popular implementations of MAP estimates work mostly under the Linear-Gaussian encoding model, which assumes additive Gaussian noise to the neural response, and are restricted to only one filter at a time.

An alternative popular approach to encode prior knowledge for estimating STRFs is to parameterize the STRF with a set of basis functions with the desired properties. In this way, model fitting becomes much more efficient as the number of parameters to be optimized is significantly reduced, and some degree of smoothness is automatically enforced. Previous studies used a raised-cosine basis for 1D retinal ganglion cell STRF in macaque monkey^7^ and Gaussian spectral or Morlet wavelet basis for 2D auditory STRF in ferrets^8^. Those basis functions are chosen to capture certain STRF properties in their corresponding studies, thus they are relatively inflexible to generalize to other settings. Besides, they are typically controlled by multiple hyperparameters, which creates a burdensome and time-consuming model selection process.

Here, we propose to use an alternative basis for receptive field inference: natural cubic splines. These are known as the smoothest possible interpolant and can be easily extended to high-dimensional settings with minimal assumptions and few hyperparameters^9^. This spline basis can be incorporated into a wide-range of existing receptive field models (Figure 1), such as the Linear-Gaussian (LG) and the Linear-Nonlinear-Poisson (LNP) model^7,10^ for single STRF estimation as well as Linear-Nonlinear-Linear-Nonlinear (LNLN) cascade models^11,12^ and spike-triggered clustering methods^12,13^ for multiple subunits estimation. When combined with proper regularization, smooth and sparse STRFs can be retrieved with very little data and in an computationally efficient manner, while their predictive performance reaches state-of-the-art level when compared with previous methods. We provide all developed methods as part of the RFE_ST_ toolbox.

**Figure 1.**
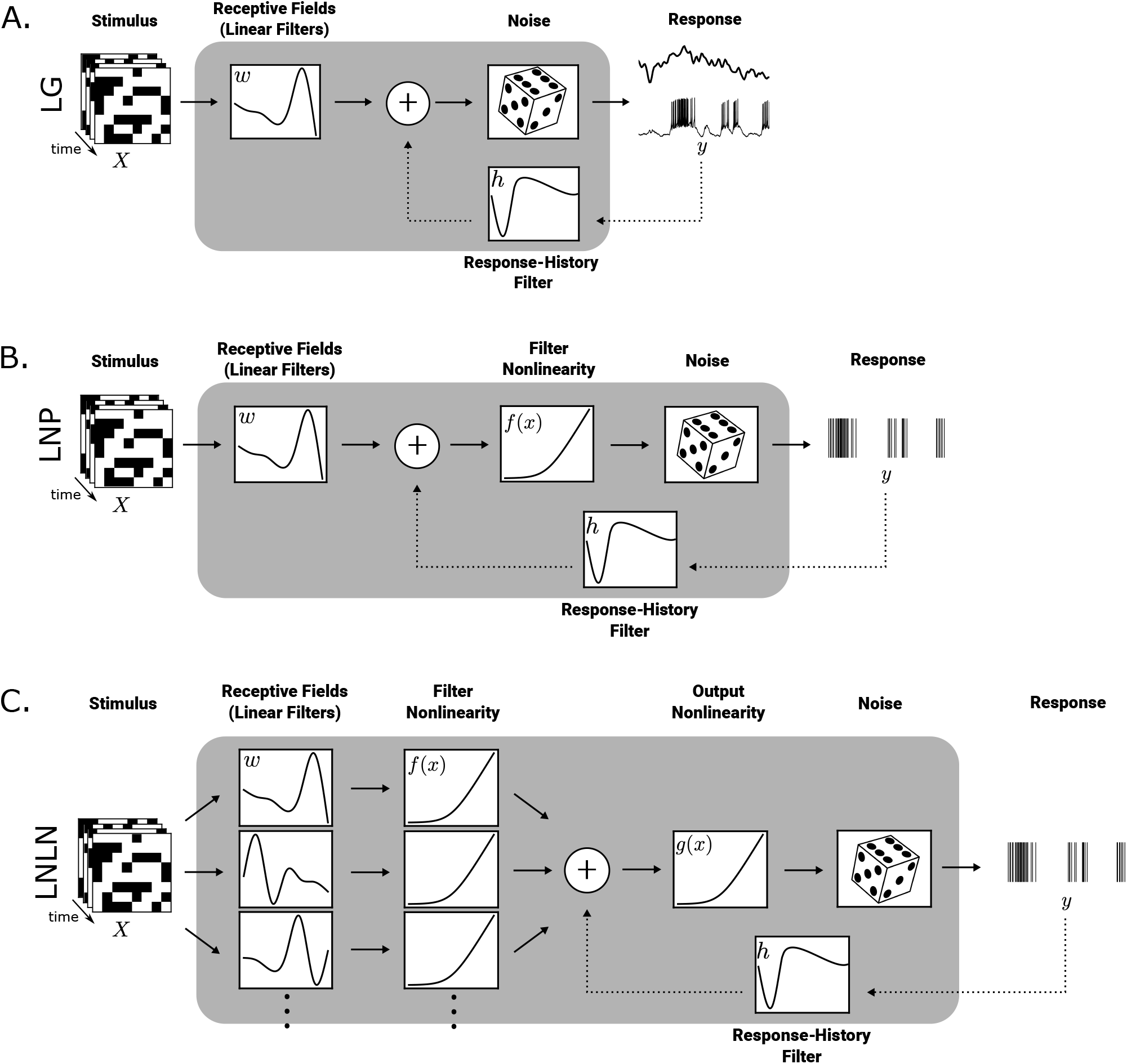
Overview of the different spatio-temporal receptive field (STRF) models used in this study. A. Linear-Gaussian (LG) encoding model. *y* = *Xw* + *ε, ε* ~ *N*(0, σ^2^), where *X* is the stimulus design matrix, *y* the response, *w* the STRF. B. Linear-Nonlinear-Poisson (LNP) model. *y* = *f* (*Xw*), *y* ~ *Poisson*, where *f* (*x*) is the filter nonlinearity. C. Linear-Nonlinear-Linear-Nonlinear-Poisson (LNLN) model. *y* = *g*(**∑** *f* (*Xw*)), *y* ~ *Poisson*, where *g* is the output nonlinearity. Optionally, an autoregressive term convolving previous response with a response-history filter can be added before the nonlinearity to improve the model predictive performance.

## 2 Results

We developed the RFEST toolbox for spline-based STRF estimation in various models, including the single-filter LG and LNP models as well as the multi-filters LNLN cascade model (Figure 1). We measured the performance of the enhanced models and compared it with their no-spline versions or other previous methods, using simulated data as well as neural recordings from the retina and the visual cortex. For the simulated benchmark data, we measured the mean squared error (MSE) between the ground truth STRF and the estimated STRFs using various methods and studied its dependence on the amount of available data. For the experimental data, we measured the correlation between the predicted and measured responses as a performance measure.

### 2.1 Estimating receptive fields from simulated LG, LNP and LNLN models

We first simulated the responses of a sensory neuron y with additive Gaussian noise using a STRF with 30 time frames and 40 pixels in space in response to white and a pink noise stimulus over 10 trials with different lengths (see Methods 4.1 for details). First, we fit LG models (Figure 1A) to these responses using various STRF estimation techniques.

We measured the MSE between the ground truth STRF and the different estimates with respect to the ratio of number of samples *n* over the number of parameters *d*, averaged over 10 different random seeds (Figure 2A). For simplicity, we omit the response history filter, but show later that it can be efficiently estimated in our framework as well (see Figure 6D).

**Figure 2.**
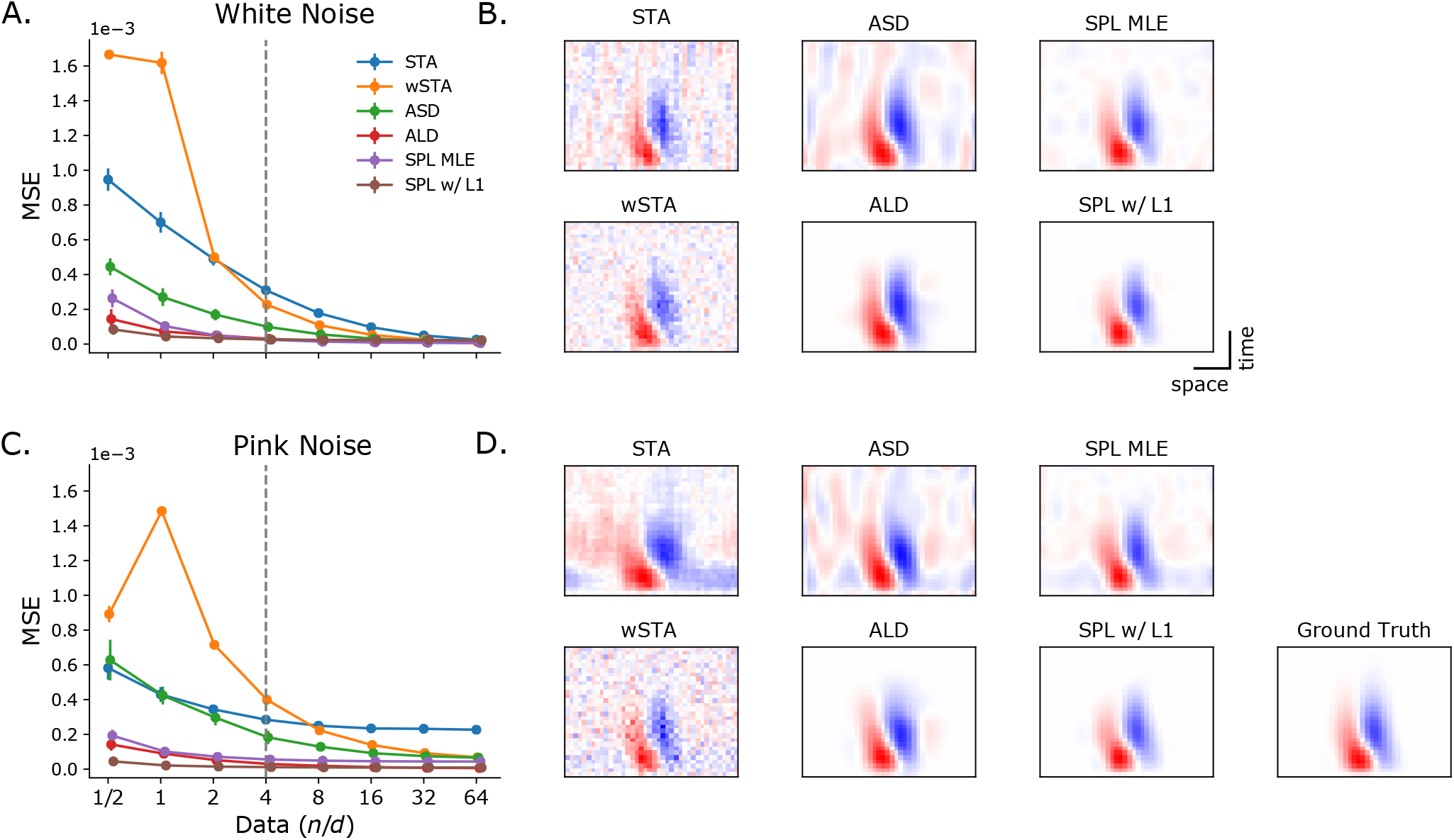
Linear-Gaussian (LG) Model. 2D simulated RF stimulated by white and pink noise in various lengths. A and C: Similarity (mean squared error, MSE) between ground Truth and estimated RFs with different length of simulated data, averaged over 10 trials with different random seeds. B and D: Example estimated RFs by different methods at n/d=4, indicated by vertical grey dash line in A and C.

As expected, the STA and wSTA approached the ground truth only in high data/feature regimes. For limited data (e.g. n/d ratio=4), the estimated STRF showed substantial background noise (Figure 2B). In the case of pink noise, the STA was in fact never able to recover the STRF due to the high correlation between pixels in the stimulus^3^ (Figure 2C-D).

We also applied more advanced techniques to estimate the STRF to the simulated data, such as automatic smoothness determination (ASD, or MAP with smoothness prior^5^) and automatic locality determination (ALD, or MAP with locality prior^6^). These recovered unbiased estimates with a relatively small amount of data, with ALD outperforming ASD alone for both white and pink noise (Figure 2). However, their runtime for estimating the optimized prior was long due to high computational complexity.

We then applied a spline-based version of the wSTA (SPL MLE) with and without L1 regularization on the basis function coefficients (see Methods). This simple change of basis functions led to an STRF estimator which performed very well, even better than ASD and ALD, especially when using L1 regularization (Figure 2). At the same time, this estimator was also computationally very efficient, as the number of coefficients to be estimated was reduced to the number of basis functions used.

The computation time between MAP-derived and spline-based GLMs was not directly comparable, as the most time-consuming part for MAP was to estimate the prior hyperparameters via evidence optimization, while spline-based GLMs need to resort to cross-validation for selecting the number of basis functions. Importantly, splines with equally-spaced knots rely only on one hyperparameter, the number of basis function *b*, also known as the degrees of freedom, for each dimension. A good strategy for selecting an optimal b would be to measure the performance of SPL MLE over a grid of possible candidates from small to large, then fit SPL w/ L1 with gradient descent to retrieve a sparse STRF.

We can still get a impression of the algorithmic efficiency of each method by profiling their most time-consuming step. Thus, we compared the time taken by the central computation of the STA (see equation 1 in Methods), wSTA (equation 2), MAP (equation 3) and SPL MLE (equation 12) for different amounts of data (Figure 3). Not suprisingly, STA was the fastest, as it had a linear time complexity (O(*n*)) and its computation time depended only on the amount of data (*n*), while wSTA and MAP were the slowest, as they were both limited by the calculation of the inverse matrix, and their computation time grew cubicly with respect to the number of coefficients (*w*) in a STRF due to the cubic time complexity (*O*(*w*^3^)) of the matrix inversion. SPL MLE, on the other hand, was less time-consuming than wSTA / MAP, even though the time complexity in general is the same. The improvement was due to the reduced number of model parameters by parameterizing the STRF coefficients with the spline bases (*b* coefficients), which resulted in a much smaller covariance matrix to invert (*O*(*b*^3^)). Thus, spline-based STRF estimators offer the best available compromise between STRF estimation quality and computation time.

**Figure 3.**
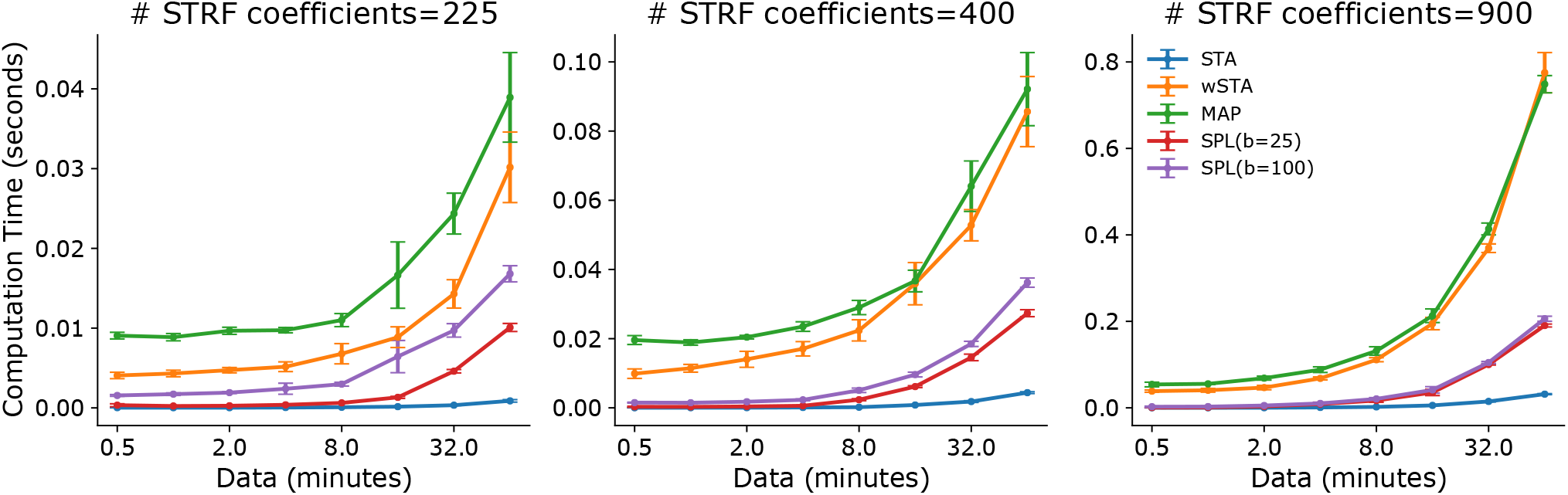
Comparison of computation time between different methods.

We next simulated responses *y* from an LNP model (Figure 1B) with Poisson noise similar to before using white and pink noise for stimulation. For simplicity, we used an exponential nonlinearity and treated it as known during the estimation procedure, but showed how it can be estimated flexibly in our framework below (Figure 6D). We then estimated the linear filter from different amounts of data using the STA, wSTA, ASD and ALD, as well as spline and no-spline LNP fitted via gradient descent with L1 regularization (referred as LNP w/ SPL and LNP, respectively). Note that the original ASD and ALD were not designed for the LNP but the LG model, nevertheless, they can still be applied to spike data as described above (with a model mismatch).

Again, we found that different methods resulted in estimates of different quality depending on the available amount of data (Figure 4). For both stimulus conditions, we found that the spline-based LNP yielded extremely data-efficient estimates of the linear filter compared to all other methods, and did so rapidly as only the coefficients for the spline basis functions needed to be optimized. In contrast, gradient-based optimization of an LNP model without spline basis, even with L1 regularization, performed only slightly better than the STA and wSTA computed under the mismatched LG model assumptions. The MAP estimators of ASD and ALD performed worse here compared to the LG case, likely because the MAP methods are optimized with respect to the mismatched cost function of the LG model. LNP w/ SPL achieved similar performance as SPL w/ L1 previously, as it is optimizing to the correct cost function, the negative log-likelihood of the LNP model.

**Figure 4.**
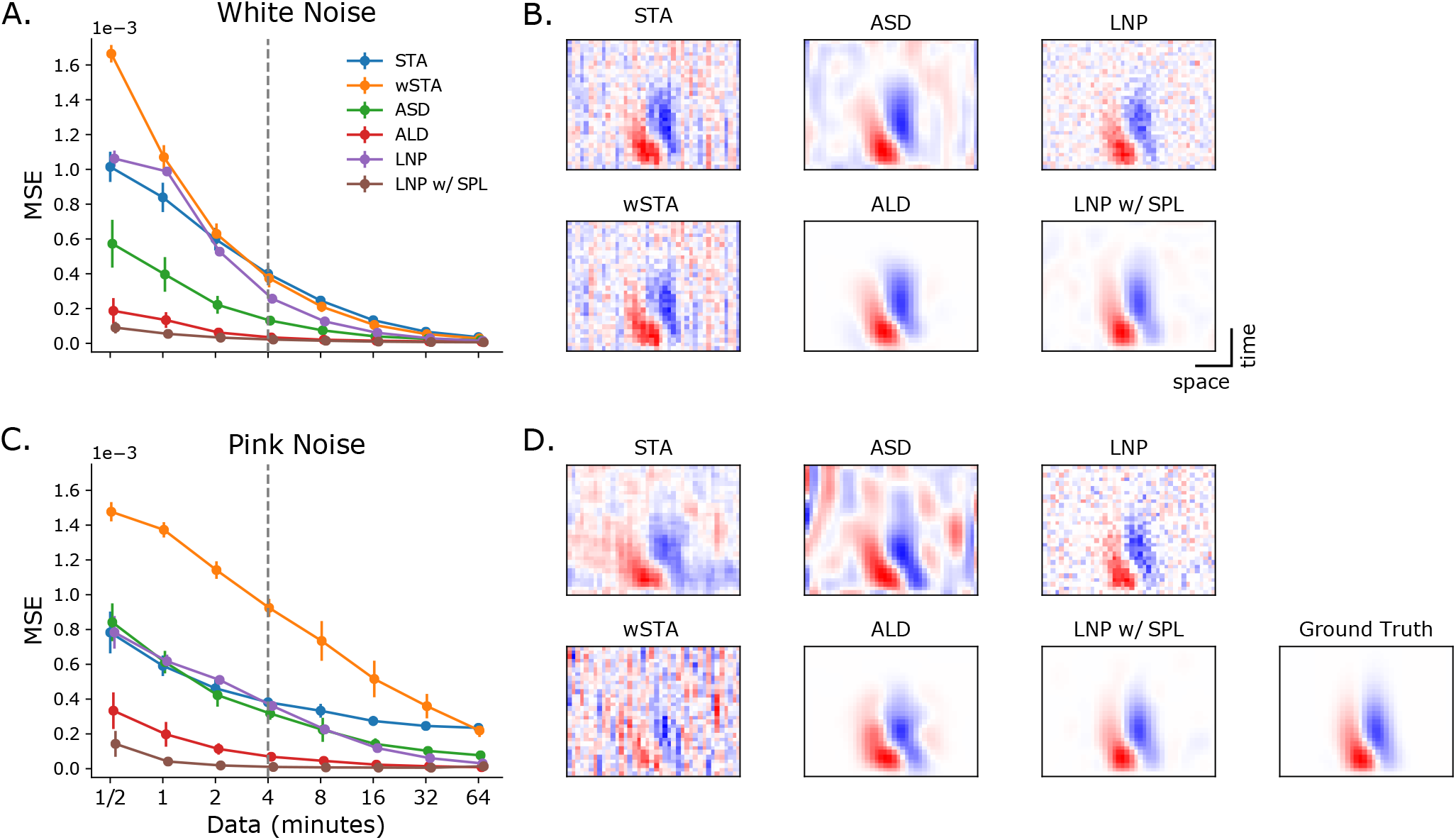
Linear-Nonlinear-Poisson (LNP) Model. 2D simulated RF stimulated by white noise and pink noise in various lengths. A and C: Details are the same as Figure 2. B and D: Example estimated RFs by different methods at data=4 minutes, indicated by vertical grey dash line in A and C.

We finally show how recently proposed subunit models (Figure 1C) can be made more efficient using a spline basis. We show this for a LNLN cascade model^11,14^ and spike-triggered clustering or non-negative matrix factorization methods^12,13^ and compare them without and with spline augmentation (see Methods 4.6). The assumption behind spike-triggered clustering methods is that each spike can be contributed by different subunits of a STRF, so that each subunit can be viewed as a mini-STA of a cluster of spikes. If this assumption holds, then a spline-approximated mini-STA should be able to serve as a better centroid for clustering, as the smoothed STA is closer to the ground truth subunit STRF. We used k-means clustering as a proof of concept in place of the previous published soft-clustering method^13^ and used a multiplicative update rule^15^ for semi-nonnegative matrix factorization (semi-NMF)^12^. Moreover, we modified the previous published semi-NMF^12^ and imposed the nonnegative constraint to the weight matrix instead of the subunit matrix, which allowed us to retrieve antagonistic STRFs.

We simulated responses *y* with Poisson noise from the model by stimulating two 20×20 pixels antagonistic center-surround subunits with different lengths of white noise stimuli. Both filter outputs first went through a exponential nonlinearity before summing together, then finally went through a softplus output nonlinearity.

We estimated subunit STRFs using k-means clustering or semi-NMF of the spike triggered ensemble, and maximum-likelihood estimation of the LNLN model directly, with or without a spline basis (Figure 5). Similar to the results from previous simulations, retrieved subunit STRFs from spline-based models were generally much more similar to the ground truth than models without spline, and required smaller amount of data to achieve similar level of performance. For example, when using a spline basis, we were able to retrieve the two subunit filters at high quality using only 4 minutes of data. With this amount of data, all other methods still essentially showed only noise with a hint of signal in the estimate of the linear filter.

**Figure 5.**
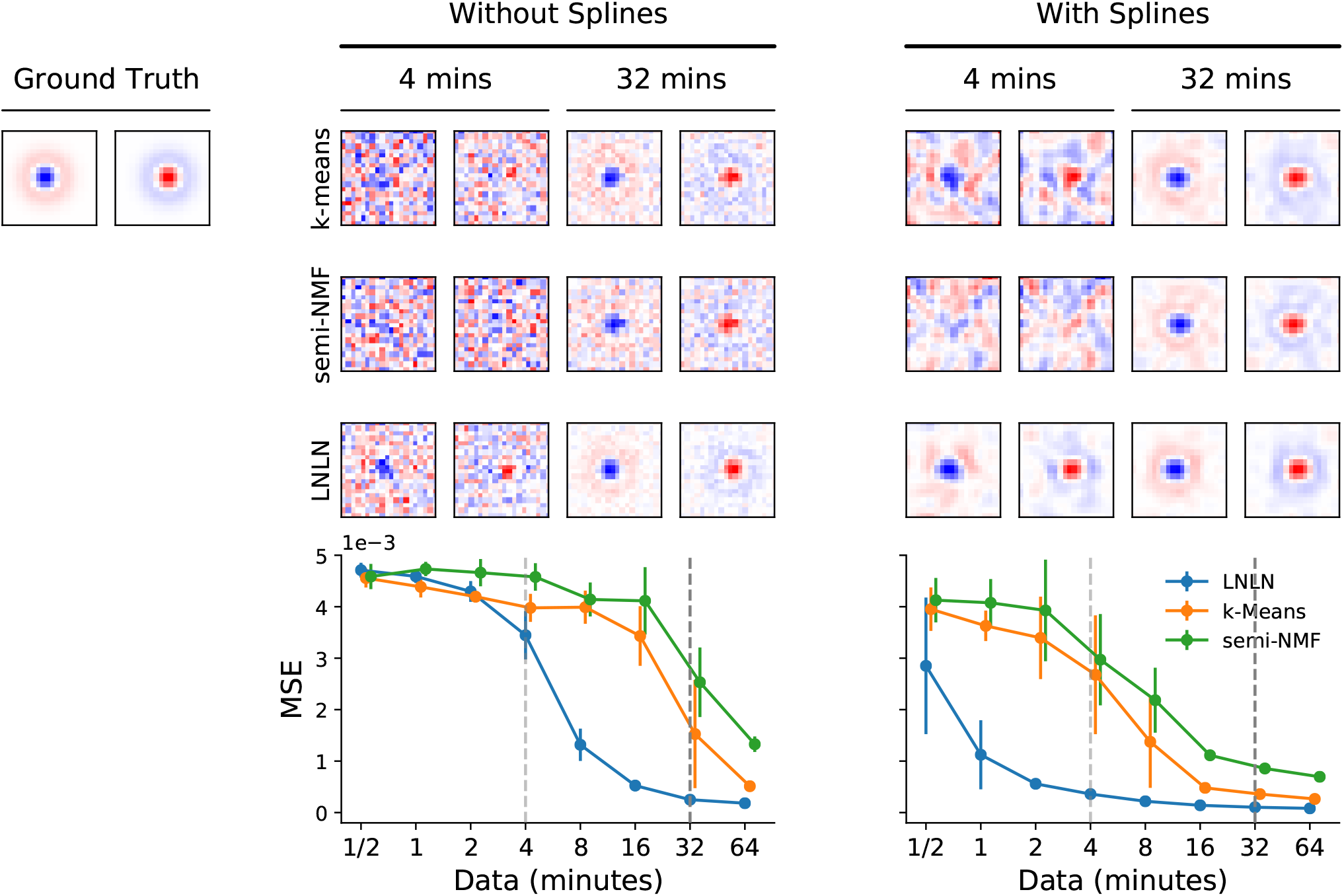
Subunit models. Two antagonistic Difference-of-Gaussian 2D RF stimulated by white noise. The subunits STRFs were estimated by k-means clustering, semi-NMF and LNLN, without and with spline. Example shown are estimated subunits by different methods at data = 4 and 32 minutes, indicated by vertical light and dark grey dash line in the bottom panel, respectively.

It is worth noting that also the computational efficiency of LNLN estimation was improved a lot by using a spline basis, as the number of coefficients to be estimated was significantly reduced, while the computation time for both spike-triggered clustering methods were actually decreased by incorporating splines, as computational steps were added to the original algorithms in those modified versions (see Methods 4.6).

### 2.2 Application to experimental data

We used gradient-based estimation for the parameters of spline-based LNP and LNLN models in three previously published experimental data sets: (1) extracellularly recorded spikes from tiger salamander retinal ganglion cells, stimulated by white noise flickering bars^14^ (Figure 6A); (2) extracellular recorded spikes from V1 neurons of macaque monkeys, stimulated by binary flicker bars^16^ (Figure 6B); (3) extracellular recorded spikes from tiger salamander retinal ganglion cells, stimulated by 3D non-repeat checkerboard white noise^12^ (Figure 6C). In addition, (4) we applied the same techniques to 2-photon calcium imaging data recorded from a mouse retinal ganglion cell soma stimulated by checkerboard white noise (Figure 6D).

**Figure 6.**
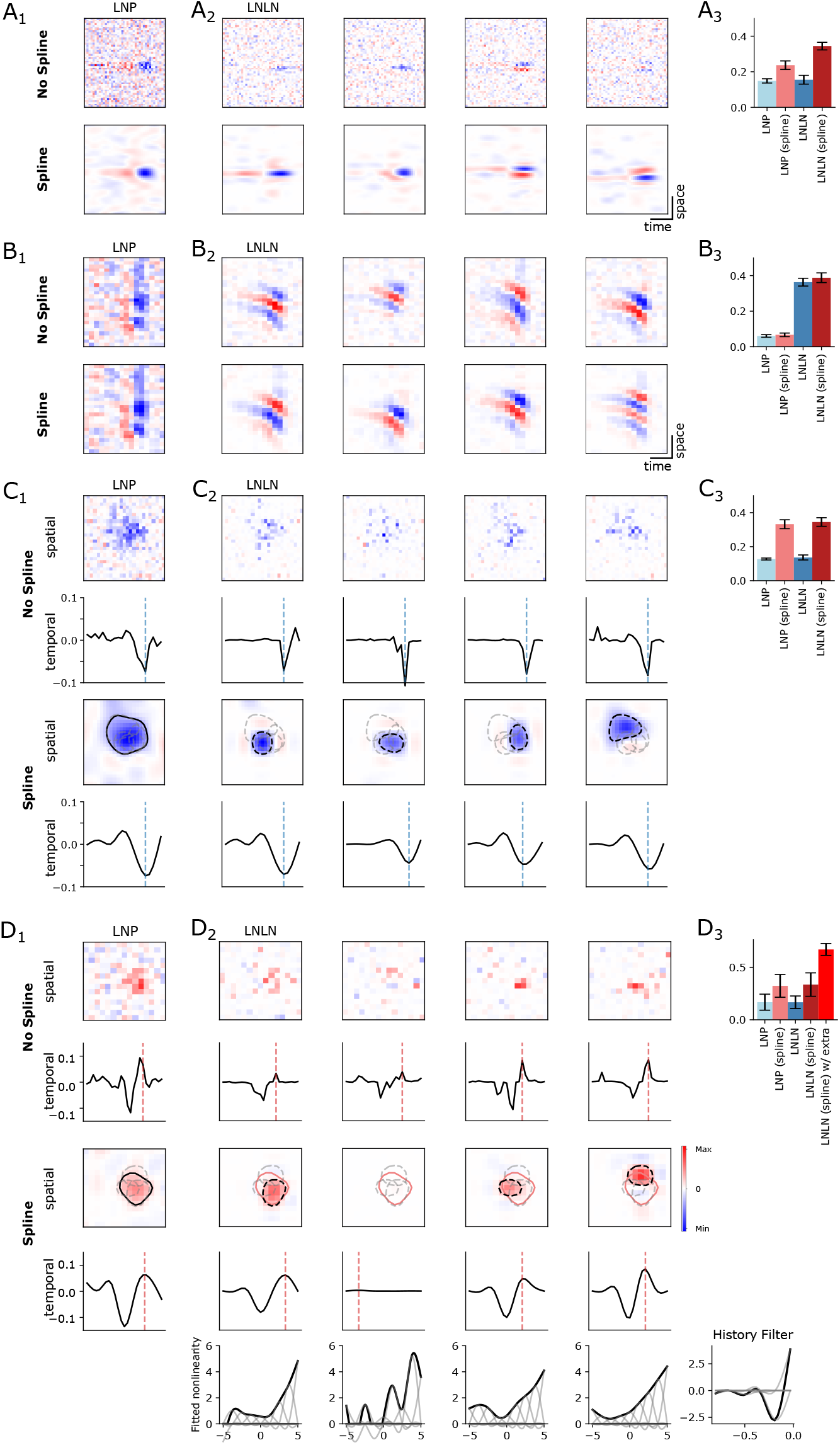
Spline-based LNP and LNLN consistently outperformed their no-spline versions on four example neurons from different data sets (see Methods 4.1 Experimental data for details). A_1_-D_1_: STRFs estimated from LNP with or without spline; A_2_-D_2_ Subunit STRFs estimated from LNLN with or without spline, where number of subunits k=4 for all cases; A_3_-D_3_: predictive performance comparison of LNP and LNLN with or without spline, measured by the averaged correlation coefficients of the test set response and the predictive ones.

For all four data sets, only the first 10 minutes of the whole recording were used for fitting the models, another 2 minutes were used for model validation and another 4 sets of 2 minutes for testing. The predictive performance was measured by the averaged correlation coefficients of the predicted and measured responses. We compared the results of spline-based models with non-spline versions. We set the number of subunits to four throughout.

For visualization purposes, STRFs were first normalized to a unit vector by dividing the norm of themselves. For 3D STRFs from data set (3) and (4), we performed singular value decomposition (SVD) to separate the spatial and temporal components. The coordinates of the extremum pixel in the spatial component were selected, and the temporal filter of the original normalized STRF is shown for those coordinates. For the spatial filter, we showed the frame of the original STRF at the extremum point of the temporal filter (indicated by a vertical dotted line). A contour was drawn on the spatial filter to illustrate the spatial extent of the STRF.

For all four example neurons shown, the spline-based LNP and LNLN consistently outperformed their no-spline versions (Figure 6). The STRFs estimated by spline-based LNP and LNLN in general were smooth and sparse with a severely limited amount of data. In contrast, STRFs estimated by no-spline models were noisy and performed comparatively worse on prediction. This is especially prominent for LNLN estimates for data set (3) and (4), as here the number of parameters to estimate exceeded the amount of available data to fit the models (Figure 6C and D). For most data sets (1), (3) and (4), the spline and non-spline version of the LNLN model improved the prediction performance only little compared to the respective LNP models (Figure 6A_3_, C_3_ and D_3_). For the complex cell in V1 (Figure 6B_3_), the LNLN model performed in general much better than the LNP model, as the subunits of V1 complex cells tend to cancel each other and result in a flat STA^16^. The spline-LNLN only improved the performance very little, possibly because of a high signal-to-noise ratio in the data, which allowed fitting the LNLN model even without splines.

For data set (3), the number of subunits might be an under-estimate as the subunits with even smaller STRFs have been shown to correspond to the STRFs of bipolar cells (Figure 6C), the neurons presynaptic of the RGCs^12,14^. This could be detected only by cross-validation with different numbers of subunits. Possibly, the number of subunits set here is an over-estimate for data set (4), because the sparsity regularization with the L1 penalty was able to push all pixels of the additional subunit to zero (Figure 6D2), while the needed ones were kept intact. While in many cases a biological interpretation of subunit STRFs may be possible in principle, the number of retrieved subunits is also influence by the quality or the signal-to-noise ratio of the data (as e.g. for data set 4), and the amount of available data. Generally speaking, the better quality and the longer the data is, the easier it is to recover the “correct” number of subunits in the model.

To further improve model prediction performance, a response-history filter (as shown in Figure 1) and flexible nonlinearity can be added to the model, both of which can also be parameterized by a 1D spline basis (Figure 6D, bottom row). With such extra flexibility, a model can better adapt to the cell-specific firing threshold and adaptation to the stimulus ensemble. The response-history filter and nonlinearity can be simultaneously optimized along with the STRF, but in practice, we found fitting them separately converged to a better result.

## 3 Discussion

Here, we showed that natural cubic splines form a good basis for STRF estimation by solving one of the main challenges in sensory neuroscience: to efficiently estimate high-dimensional STRFs with limited data. Using this basis function set, we showed that inference for single filter LN-models with different noise models as well as LNLN cascade models can be efficiently performed from both spiking as well as two-photon imaging data, even if the signal-to-noise ratio is low (and see also how spline-based methods compare to previous methods in Table 1). We provide an implementation of the proposed methods in the RFEST-package for Python.

**Table 1.** Model comparison. Here, *n* is the number of samples, *w* the number of STRF coefficients, *b* the number of spline basis coefficients and *k* the number of subunits. In most cases, *b* ≪ *w*.

| Methods | Stimuli Type | Noise Distribution | Data needed | Complexity | Multiple Filters |
| --- | --- | --- | --- | --- | --- |
| STA | White Noise only | Gaussian | a lot | $O(n)$ | No |
| wSTA | Any | Gaussian | a lot | $O(w^3)$ | No |
| MAP | Any | Gaussian | little | $O(w^3)$ | No |
| LNLN | Any | Poisson | a lot | $O((wk)^3)$ | Yes |
| SPL LG | Any | Gaussian | little | $O(b^3)$ | No |
| SPL LNP | Any | Poisson | little | $O(b^3)$ | No |
| SPL LNLN | Any | Poisson | little | $O((bk)^3)$ | Yes |

### 3.1 Choice of basis functions

We chose the natural cubic spline basis to represent the linear filters as default in our toolbox, because among many other spline bases such as cyclic cubic splines, B-splines, or thin-plate splines, natural cubic splines have been shown to be the smoothest possible interpolant through any set of data and theoretically near-optimal to approximate any true underlying functions closely^9^. Conveniently, assuming all knots are spaced equally in each dimension, natural cubic splines only require setting one hyperparameter: the degree of freedom, or the number of basis functions. This is unlike the raised cosine basis used previously^7^, which is governed by multiple hyperparameters. Also, natural cubic splines can be easily extended to high-dimensional setting, also known as tensor product bases, by simply taking the row-wise Kronecker product of the basis matrix along each dimensions (see Method 4.4), making them uniquely suitable for estimating spatio-temporal receptive fields.

Through extensive validation with simulated and real experimental data sets, we showed that STRFs estimated with this basis can not only be more accurate than previous state-of-the art methods but the estimation procedure was also computationally much more efficient in terms of both time and memory. The estimated STRFs were smoothed automatically, as they were the linear combination of a set of smooth basis functions. Further, sparsity and localization could often be achieved by adding a L1 regularization term to the loss function, which pushed the coefficient for the less relevant basis functions to zeros. Moreover, by parameterzing STRFs with basis functions, the number of coefficient could be significantly reduced, hence model fitting became much more computationally efficient, and the amount of data needed for the optimization to converge was also significantly less.

### 3.2 Other approaches to efficient estimation of receptive fields

Similar to our approach, fast automatic smoothness determination (fASD)^17^ has been suggested to speed up the process of evidence optimization in ASD. Because the smoothness prior covariance can be represented in frequency coordinates and many high frequency coefficients of the Fourier-transformed filter tend to be small, the size of the prior covariance matrix can be reduced, leading to a speed up in the optimization process. However, as the efficiency of fASD is controlled completely by the initial frequency truncation, and the level of frequency to be truncated have to be fixed before the evidence optimization starts, one needs to resort back to grid search and cross-validation for choosing an good starting point, making the automatic hyperparameters selection via evidence optimization less automatic and comparable to the need to find the optimal degrees of freedom for the spline basis. Importantly, fASD is so far only applicable within the framework of linear model with Gaussian additive noise, while our approach is ready to use in nonlinear model with non-Gaussian noise, as shown for an LNLN model.

Regarding the inference for multiple STRFs in LNLN-cascade models, previous work has attempted to provide efficient algorithms by first retrieving subunit STRFS through spike-triggered clustering methods such as soft-clustering^13^ or semi-NMF^12^. These were subsequently used to estimate other model parameters via gradient descent while holding the subunit STRFs fixed. Utilising only the collection of stimuli that elicit a spike, these approaches have advantages in terms of the computational cost in time and memory compared to the direct fitting of the complete model using gradient descent^11,14^.

The ease of using these algorithms relying on the spike-triggered ensemble under white noise stimulation is also the disadvantage of these approaches – they can only be applied to spike data under white noise stimulation. In many cases, experiments today record more diverse measurements from neurons, such as two-photon calcium imaging^18,19^ or synaptic current recording^20^ under more diverse types of stimulation, such as correlated noise and natural stimuli^21^. In those cases, spike-triggered analysis can not be used directly without modification, as the spike-triggered average is no longer the maximum likelihood estimate^3^. In contrast, our method does not rely on the Gaussianity of the stimulus or the discreteness of the response, and in principle can be generalized to other stimuli distribution and other response types.

In addition, current spike-triggered clustering methods rely on temporally collapsing the 3D stimuli to further reduce the model complexity, assuming all subunits share the same temporal response kernel. This limits the usability of these methods to some very specific cell types, and will not account for the spatio-temporal inseparable computation implemented by neurons receiving inputs from multiple sources, such as bi-stratified On-Off direction selective cell in the retina^22^. In this case, a more general approach such as our method is preferred.

### 3.3 Further extensions

Here, we only showcased how to estimate STRFs using natural cubic spline in a set of popular models with standard gradient descent with L1 regularization. This simple spline add-on can be directly incorporated into other previous models^11,14^ with different constraints, regularization and optimization techniques, as it only changed the linear step of the models without modifying other parts.

Furthermore, we implemented a spline-based response-history filter (as shown in Figure1 and Figure 6D) as a useful option for improving model predictive performance. Still, even with the response-history filter, a STRF model can only account partially for the total variability of the neuronal responses, meaning such variability might be contributed from other sources. Future extension of the current GLM for STRFs can consider incorporating more diverse sources of behavioral data^23^ of the experimental animal under experimental or naturalistic settings as 1D time-dependent filters.

## 4 Methods

### 4.1 Data sets

#### Simulated data

For the simulated data shown in Figure 2 and 4, we used a STRF with 30 time frames and 40 pixels in space (as shown in both Figure as ‘‘ground truth”) and generated responses with two types of flicker bar stimuli: Gaussian white noise and pink noise. We used a fixed exponential nonlinearity for LNP models. For Figure 5, we used two 20×20 pixels antagonistic center-surround STRF and only 2D Gaussian white noise was used as stimulus. We used a fixed exponential function for the filter nonlinearity and a softplus function for the output nonlinearity.

For all simulations, we measured the similarity of the ground truth STRF and the model estimators by computing the MSE between the two, after dividing by the norm of themselves.

For Figure 2, the similarity is measured at various length of stimulus (*n*_samples_), which were scaled by different factors (factor ∈ 0.5, 1, 2, 4, 8, 16, 32, 64) respect to the number of coefficients of the STRF. For Figure 4 and 5, the similarity is measured at various length of time (time ∈ 0.5, 1, 2, 4, 8, 16, 32, 64 minutes)), which is equal to time = *n*_samples_ × **Δ**, where **Δ** is the time bin size and took the value of 0.033 s.

Error bars in Figure 2–5 were obtained by averaging 10 trials of simulations with different random seeds.

#### Experimental data

We used the following previously published data sets for Figure 6A-C:

1. Extracellularly recorded spikes from tiger salamander retinal ganglion cells^14^, available to download at https://github.com/baccuslab/inferring-hidden-structure-retinal-circuits;
2. Extracellular recorded spikes from V1 neurons of macaque monkeys^16^, available on request from the authors;
3. Extracellular recorded spikes from tiger salamander retinal ganglion cells^12^, available to download at https://gin.g-node.org/gollischlab/Liu_etal_2017_RGC_spiketrains_for_STNMF

And the detail of the fourth data set for Figure 6D is described below.

### 4.2 Experimental procedures for 2-photon calcium imaging

#### Animals and tissue preparation

One wild-type mouse (C57Bl/6J, JAX 000664) housed under a standard 12 hour day/night rhythm was used in this study. Before experiments, the mouse were dark adapted for at least 2 hours. Then, the animal was anaesthetized with isoflurane (Baxter) and killed with cervical dislocation. Eyes were quickly enucleated in carboxygenated (95% O_2_, 5% CO_2_) artificial cerebral spinal fluid (ACSF) solution containing (in mM): 125 NaCl, 2.5 KCl, 2 CaCl_2_, 1 MgCl_2_, 1.25 NaH_2_PO_4_, 26 NaHCO_3_, 20 glucose, and 0.5 L-glutamine (pH 7.4). After carefully removed the cornea, sclera and vitreous body, the retina was flattened on an Anodisc (0.2 *μ*M pore size, GE Healthcare) with the RGC side facing up and electroporated (see below). Then, the retina was transferred to the recording chamber of the microscope. To visualize the blood vessels and the damaged cells, we added 5 *μ*M sulforhodamine-101 (SR101, Invitrogen) into 2l of ACSF solution. All experimental procedures were carried out under very dim red light. All animal procedures were approved by the governmental review board (Regierungspräsidium Tübingen, Baden-Württemberg, Konrad-Adenauer-Str. 20, 72072 Tübingen, Germany) and performed according to the laws governing animal experimentation issued by the German Government.

#### Electroporation

For two-photon Ca^2+^ imaging, Oregon-Green BAPTA-1 (OGB-1, hexapotassium salt; Life Technologies) was bulk electroporated^24^. After the retina was flattened on an Anodisc and placed between two 4-mm horizontal plate electrodes (CUY700P4E/L, Nepagene/Xceltis), 10 μl OGB-1 (5 mM) diluted in ACSF solution was suspended on the upper electrode. Then the upper electrode was lowered onto the retina and applied 10 pulses (9.2 V, 100 ms pulse width, at 1 Hz) from a pulse generator/wide-band amplifier combination (TGP110 and WA301, Thurlby handar/Farnell). For complete recovery of the electroporated retina, we started recordings 30 min after electroporation.

#### Two photon imaging and light stimulation

A MOM-type two-photon microscope (designed by W. Denk, MPI, Martinsried; purchased from Sutter Instruments/Science Products) as described previously^25,26^ was used for this study. Briefly, the system was equipped with a mode-locked Ti:Sapphire laser (MaiTai-HP DeepSee, Newport Spectra-Physics, Darmstadt, Germany), green and red fluorescence detection channels for OGB-1 (HQ 510/84, AHF, Tübingen, Germany) and SR101 (HQ 630/60, AHF), and a water immersion objective (16x/0.80W, DIC N2, **∞**/0 WD 3.0, Nikon). For Ca^2+^ imaging, we tuned the laser to 927 nm, and used a custom-made software (ScanM, by M. Müller, MPI, Martinsried, and T.E.) running under IGOR Pro 6.3 for Windows (Wavemetrics). Time-elapsed Ca^2+^ signals were recorded with 64× 16 pixel image sequences (31.25 Hz).

Light stimuli were projected through the objective lens^26^. A digital light processing (DLP) LightCrafter 4500 (Texas Instruments, Dallas, TX) coupled with external unit with light-emitting diodes (LEDs) – “green” (576 nm) and UV (387 nm) that match the spectral sensitivities of mouse M- and S-opsins (for details, see^18,27^) was used in this study. Both LEDs were intensity-calibrated to range from 0.1 · 10^3^ (“black” background) to 20.0 · 10^3^ (“white” full field) photoisomerisations P*s^−1^ per cone. The light stimulus was centred before every experiment, ensuring that its centre corresponded to the centre of the microscope’s scan field. For all experiments, the tissue was kept at a constant mean stimulator intensity level for ≥ 15 s after the laser scanning started and before light stimuli were presented.

Binary dense noise (20× 15 matrix of 30×30 *μ*M pixels; each pixel displayed an independent, balanced random sequence at 5 Hz for 20 minutes) were generated and presented using the Python-based software package QDSpy (https://github.com/eulerlab/QDSpy).

#### Data pre-processing

Ca^2+^ imaging data were pre-processed using custom-written scripts in IGOR Pro. Regions-of-interest (ROIs) were manually defined depending on the outline of different soma. For each ROI, Ca^2+^ traces were extracted using the image analysis toolbox SARFIA for IGOR Pro^28^. A stimulus time marker embedded in the recorded data served to align the traces relative to the light stimulus at a temporal resolution of 2 ms. All stimulus-aligned traces together with the relative ROI position were exported for further analysis.

### 4.3 Previous methods

#### Spike-triggered Average (STA)

The STA was computed as the average of all stimuli that proceeded a spike^1^:

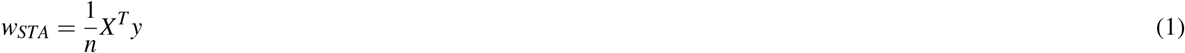

where *X* is the stimulus design matrix, *y* the response, *n* can be the number of spikes or the number of samples (rows) in *X* and *y* for non-spike data.

#### Whitened spike-triggered average (wSTA)

The maximum-likelihood estimator under a Gaussian noise model, or whitened-STA (wSTA), was computed by multiplying the STA with the inverse of the stimulus auto-correlation matrix:

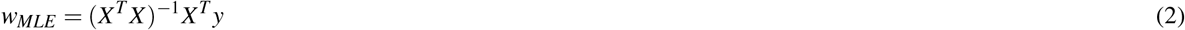

But in practice and in our code base, we did not compute the matrix inverse explicitly but compute the least squared solution instead.

#### Maximum-a-posteriori estimator (MAP)

Similar, the MAP under a Gaussian noise model was computed by adding the inverse of the prior covariance to the stimulus auto-correlation matrix:

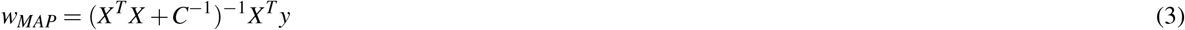

where *C* is a prior covariance matrix of choice, e.g. corresponding to a smoothness or sparseness assumption^5,6^. Its hyperparameters can be optimized by minimizing the negative log-evidence^5,6^:

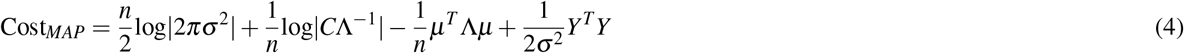

where σ is variance of the additive Gaussian noise, *μ* and **Λ** are posterior mean and covariance, respectively:

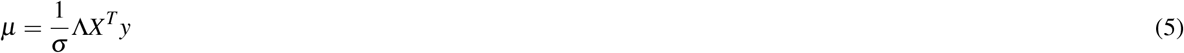

and

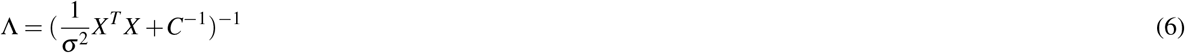

#### Smoothness prior covariance matrix for ASD

The 1D ASD prior covariance matrix is formulated as following^5^:

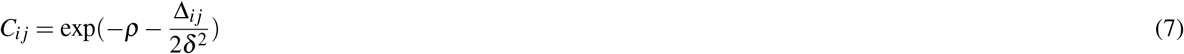

where Δ*ij* is the squared distance between STRF coefficients *w_i_* and *w_j_, ρ* controls the scale and *δ* controls the smoothness.

#### Locality prior covariance matrix for ALD

The 1D ALD prior covariance matrix is formulated as following^6^:

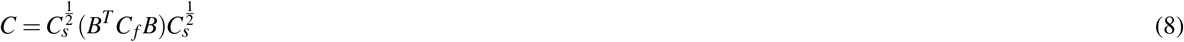

where *B* is an orthogonal basis matrix for the 1D discrete Fourier transform, *C_s_* and *C_f_* are the diagonal ALD prior covariance in space and frequency, respectively, which can be constructed as:

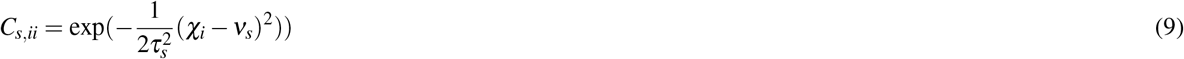

where *τ_s_* is scalar values control the shape and extent of the local region, *χ* is the STRF coefficient coordinates in spacetime and *ν_s_* is center coordinate of the STRF. And

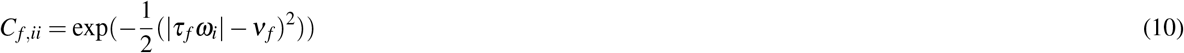

where *τ_s_* is scalar values control the scale of smoothness in frequency, *ω* is of STRF coefficient coordinates in frequency and *ν_f_* is center coordinate of the STRF in Fourier domain.

High-dimensional prior covariance matrix can be approximated by taking the Kronnecker product (⨂) of the 1D covariance matrix in each dimension.

### 4.4 Estimating STRFs with Natural Cubic Regression Splines

To generate a natural cubic spline matrix (*S*), we used the implementation from PATSY^29^, which is a Python equivalent of the original implementation in R package MCGV^9^. The only hyperparameter needed to generate a 1D spline basis matrix is the number of basis functions, or the degrees of freedom (*df*), as we assume that all knots are equally-spaced.

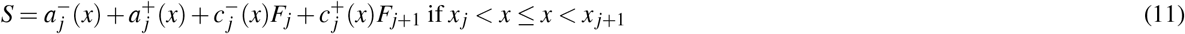

where *S* is the spline basis matrix, *x* is knot locations, *F*^−1^ = *B*^−1^ *D, F* = (0, *F*^−1^, 0) and 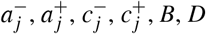 are defined in Table 2.

**Table 2.**
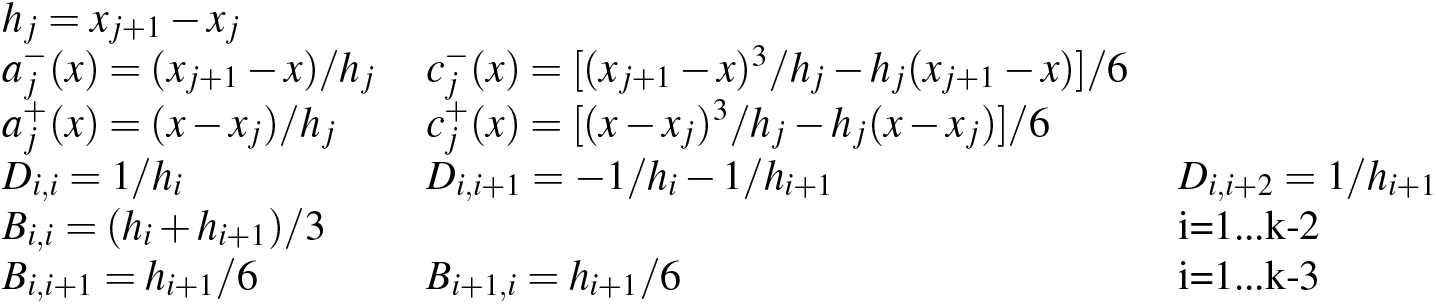
Basis functions for natural cubic spline.

An illustration of natural cubic splines can be seen in Figure 7. A spline-based STRF under a Gaussian noise model can be fitted in the similar manner as the wSTA:

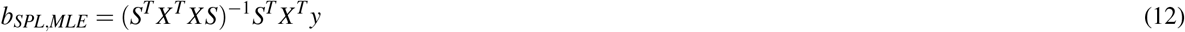

where *b* is the spline coefficients, and the corresponding RF is *w_SPL,MLE_* = *Sb_SPL,MLE_* = *S*(*S^T^X^T^XS*)^−1^*S^T^X^T^y*.

**Figure 7.**
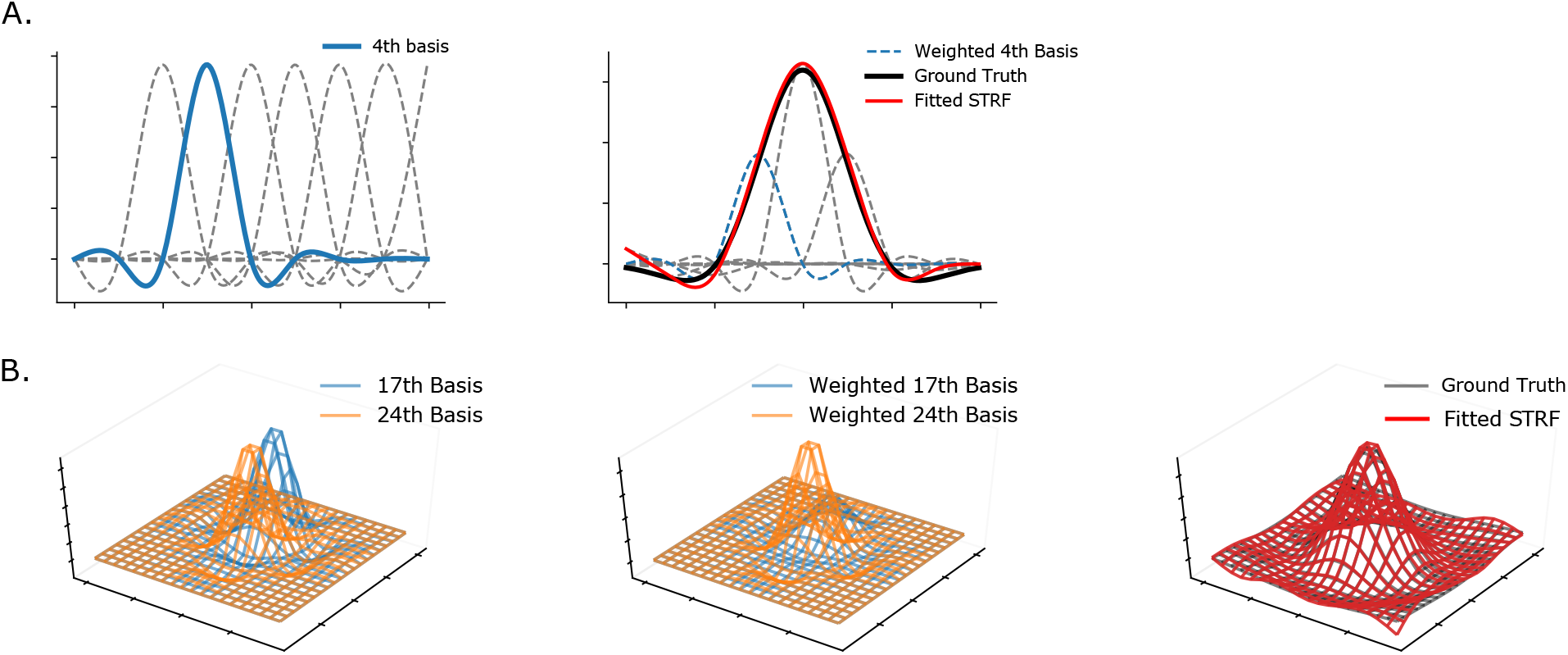
Illustration of the Natural cubic spline basis. A. 1D natural cubic spline. Left: a set of cubic spline basis, with the 4rd basis highlighted. Right: how basis functions are combined and fitted to a simulated 1D RF as a smooth curve. B. 2D natural cubic spline. Left: Two selected bases of 2D natural cubic spline. Middle: Weighted bases after fitted to a simulated 2D RF. Right: An example of a fitted 2D STRF overlaid on the Ground Truth.

The spline matrix can be extended to high-dimensions, which is also called tensor product smooth^9^, by simply taking the Kronnecker product (⨂) of the basis functions in each dimension:

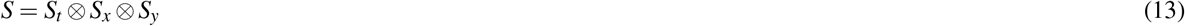

### 4.5 Spline-based GLMs with gradient descent

The spline coefficients can also be fitted by gradient descent with respect to the cost function defined in each model with added regularization through L1 and L2 penalties.

For the LG model, the cost function is the mean square error:

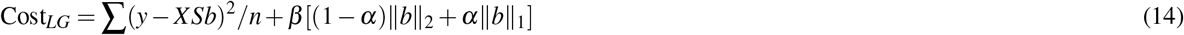

where *α* is the balancing weight for L1 and L2 regularizers, *β* is the overall weight of regularization.

For LNP and LNLN model, the cost function is the negative log-likelihood:

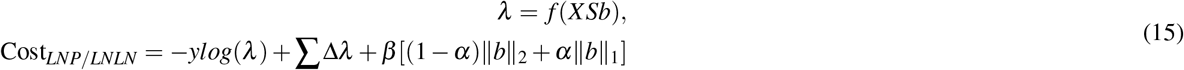

where *λ* is the conditional intensity or the instantaneous firing rate, **Δ** is the size of time bin, and *f* (.) is a nonlinearity applied to the filter output, which can be fixed as a softplus or exponential nonlinearity or be parameterized by a set of spline basis (*ϕ*):

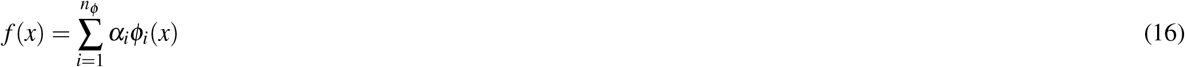

where *α* is basis weight.

We used JAX^30^ for automatic differentiation and parameters optimization.

### 4.6 Spline-based spike-triggered clustering methods

Both k-means clustering and semi-NMF can be formulated as a matrix factorization problem:

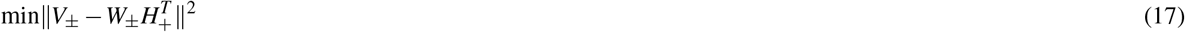

where *V* is the spike-triggered ensemble, *W* is clustering centroids matrix (or subunit matrix), and *H* is the clustering label (or subunit weight matrix). A spline basis can be easily incorporated into k-Means clustering by assuming that each centroid can be approximated by the same spline basis matrix: *W_SPL,centroid_* = *SB*, where *B* is the spline coefficient matrix for each subunit, and can be computed as the least-squares solution to the linear matrix equation *B* = *S*^†^W_centroid_, where † is pseudo-inverse. For semi-NMF, splines can also be combined with the update rules^15^:

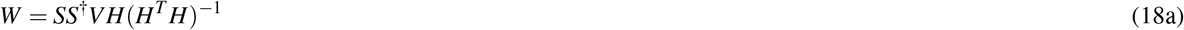

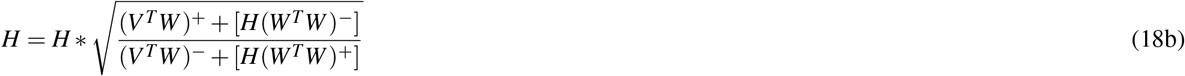

where * is point-wise multiplication, *A*^+^ = (| *A* | +A)/2 and *A*^−^ = (| *A* | – A)/2.

### 4.7 Comparison of Computation Time

For comparison of computation time in Figure 3, we used simulated 2D RFs of different sizes (15×15, 20×20 and 30×30 pixels, respectively) and white noise stimuli. Simulations were carried out at different length of time in the same manner as Figure 4 and 5. The number of basis for SPL MLE were fixed as 5 and 10 along each dimension in all three cases. Measurements of computation time were averaged over 10 repetitions and performed on a 2019 16-inch MacBook Pro with a 2.3 GHz 8-core Intel Core i9 and 16 GB of RAM.

## 4.8 Data availability

All data used, except the data set (2) shown in Figure 6B, are available in Github (https://github.com/huangziwei/data_RFEst). Data set (2) is available on request from the authors of^16^.

## 4.9 Code availability

All methods (spline-based GLMs, evidence optimization and spike-clustering methods) were implemented in Python3 and available in GitHub (https://github.com/berenslab/RFEst). And Jupyter notebooks for all figures are available also in Github (https://github.com/huangziwei/notebooks_RFEst)

## Acknowledgments

We thank the authors of^14^,^12^ and^16^ for making their data available. Research was funded by the Deutsche Forschungsgemeinschaft through a Heisenberg Professorship (BE5601/4-1 and BE5601/8-1, PB), the Collaborative Research Center 1233 “Robust Vision” (ref number 276693517) as well as individual research grants (EU 42/10-1, BE5601/6-1), the German Ministry of Education and Research through the Bernstein Award (01GQ1601, PB). PB is a member of the Excellence Cluster 2064 “Machine Learning — New Perspectives for Science” (ref number 390727645) and the Tübingen AI Center (FKZ: 01IS18039A) and funds from the Ministry of Science, Research and Arts of the State of Baden-Württemberg.

